# Spatially Enhanced Processing for Valuable Objects in Prefrontal Cortex Neurons During Efficient Search

**DOI:** 10.1101/2024.04.08.588532

**Authors:** Kiomars Sharifi, Mojtaba Abbaszadeh, Ali Ghazizadeh

**Affiliations:** Bio-intelligence Research Unit, Sharif Brain Center, Electrical Engineering Department, Sharif University of Technology, Tehran, Iran; School of Cognitive Sciences, Institute for Research in Fundamental Sciences (IPM), Tehran, Iran

**Keywords:** visual search efficiency, valuable object pop-out, prefrontal cortex, long-term value, receptive field, plasticity, macaque monkey

## Abstract

It is recently shown that objects with long-term reward associations can be efficiently located during visual search. The neural mechanism for valuable object pop-out is unknown. In this work, we recorded neuronal responses in the ventrolateral prefrontal cortex (vlPFC) with known roles in visual search and reward processing in macaques while monkeys engaged in efficient vs inefficient visual search for high-value fractal objects (targets). Behavioral results and modeling using multi-alternative attention-modulated drift-diffusion (MADD) indicated that efficient search was concurrent with enhanced processing for peripheral objects. Notably, neural results showed response amplification and receptive field widening to peripherally presented targets in vlPFC during visual search. Both neural effects predict higher target detection and were found to be correlated with it. Our results suggest that value-driven efficient search independent of low-level visual features arises from reward-induced spatial processing enhancement of peripheral valuable objects.

## Introduction

Low-level guiding features such as color, size, and orientation facilitate quick detection of targets in visual search through a phenomenon known as visual pop-out^1,2^. Such visual pop-out is thought to arise from the brain’s ability to simultaneously process certain low-level features in parallel across visual space in early visual areas^3–5^. However, recent studies reveal that high-level features such as value can capture attention toward peripherally presented objects^6–8^. We refer to this phenomenon as value pop-out, where high-value objects, among many distractions, automatically attract attention. Indeed, we have previously shown that this attention-capturing effect allows targets with long reward associations to be identified efficiently in visual search^9^. Importantly, such value-driven efficient search happens for complex fractal patterns and has the capacity to support pop-outs for many objects. However, unlike visual pop-out, pop-out of valuable arbitrary shapes cannot be easily explained by extant frameworks such as feature integration theory^9^ or guided search^1^ and remains to be addressed.

Recent studies have shown that cortical regions, particularly the ventrolateral prefrontal cortex (vlPFC)^10^ and subcortical areas, such as substantia nigra reticulata (SNr)^12^, show enhanced differentiation of target present and absent conditions in efficient visual search for valuable targets. Among these regions, the vlPFC is known to have a more localized receptive field (RF) that can signal both the presence of the target and its location ^8,11^. Thus, we hypothesized that the value pop-out could arise from the reward-driven enhancement of the visual processing for valuable objects in vlPFC. Such enhancement would be most beneficial for objects not yet foveated as it can aid in guiding future saccades to acquire search targets.

To address this issue, we analyzed data from an experiment in which macaque monkeys performed both efficient and inefficient value-driven visual searches while recording vlPFC neural responses^10^. Analysis of visual search behavior using a drift-diffusion model and assessing the location-dependence of neural responses in vlPFC revealed that consistent with our hypothesis, long-term reward-learning induced enhanced spatial processing of peripheral objects in ways that were predicted to allow for faster detection of a valuable object in peripheral vision.

## Results

Acute neural recordings from vlPFC (areas 8Av, 46v, and 45a) were conducted in two monkeys (monkeys S and H with 240 and 356 neurons in the left and right hemispheres, respectively). Monkeys learned to associate abstract fractal objects with either large or small juice amounts as rewards in a biased value training task (Fig 1A-B). The large number of fractals used and their arbitrary assignment in high or low-value groups ensured that the subsequent search task could not be solved by low-level guiding singletons such as unique colors or shapes. Each value training session involved a set of eight fractals, half associated with small rewards and the other half with large rewards (bad and good objects, respectively). These object sets underwent biased value training for either one day or +5 days before being used in the visual search task (Fig 1C).

**Figure 1.**
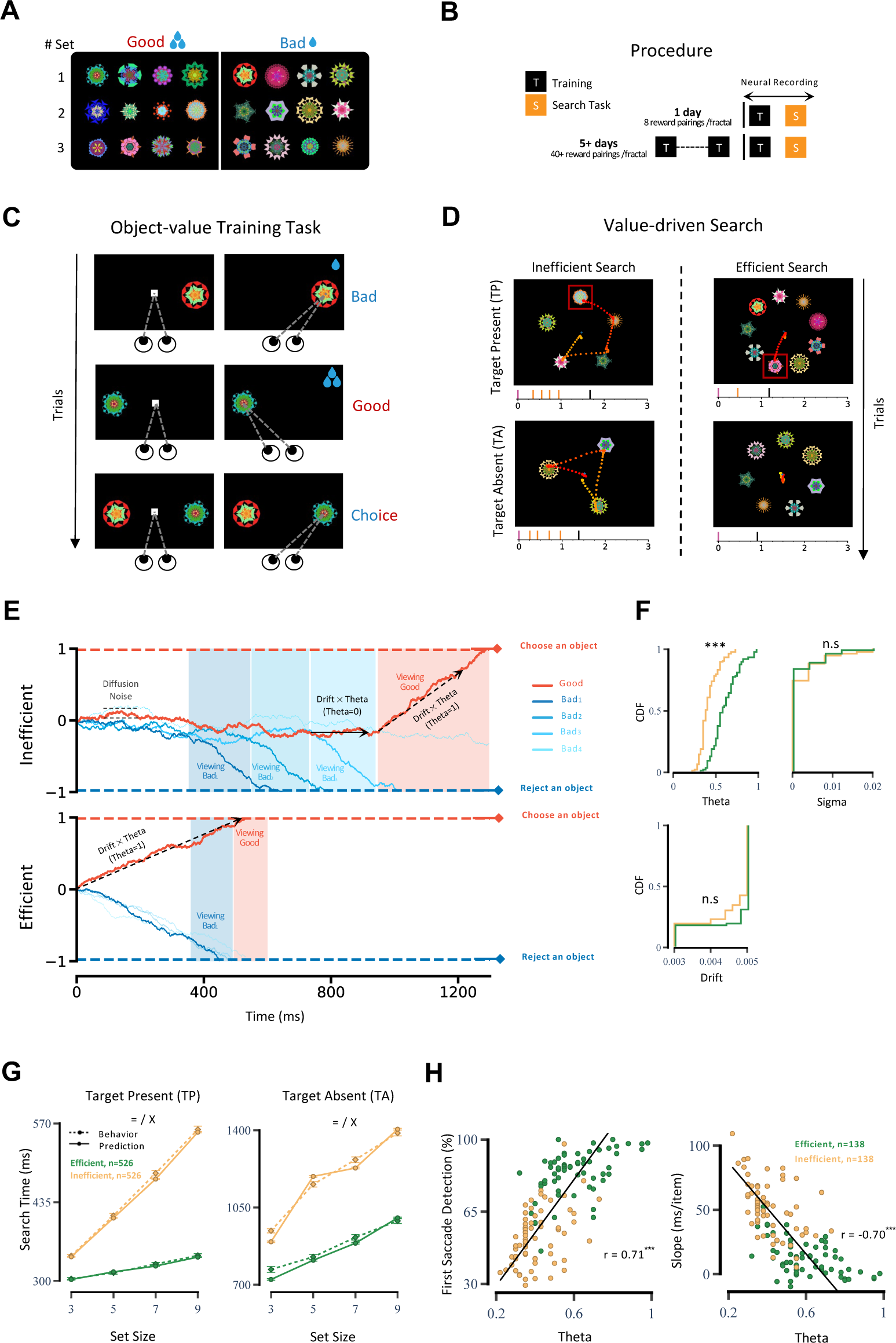
Tasks paradigm, behavioral Data, and MADD Model. (A) Example fractal stimuli used in value training and search tasks. Fractals were trained in sets of eight: four associated with a large (good) and four with a small juice reward (bad). Each search session used three sets (24 fractals, 12 good). (B) Two different groups of sets were trained for one day and 5+ days in the object-value training task. The variable training sessions were known to create a difference in the efficiency of the search tested subsequently. (C) Object-value training tasks consisted of force (80%) and choice (20%) trials. After central fixation, an object appeared in the periphery (9.5°). After the monkey made a saccade and held a gaze on the object, the corresponding large or small reward was delivered (biased reward training). (D) Value-driven search tasks consisted of target-present (TP) trials with one good object among bad objects or target-absent (TA) trials with all bad objects. The number of objects shown was variable (set size 3,5,7,9). Monkeys had 3sec for each search trial and were free to make as many saccades. Holding gaze on an object meant choosing it and was followed by a corresponding reward. Staying at the center or returning back to it meant rejecting the trial (see methods). Example trials and eye-traces for inefficient and efficient searches are color-coded by time (from orange to red). Tick marks at the bottom of trials show the timings of saccades (orange) and reward (black) relative to the display onset (purple). Note improvement of target selection in TP and trial rejection in TA in efficient search. (E) <u>M</u>ulti-alternative <u>a</u>ttention-modulated <u>d</u>rift-<u>d</u>iffusion model (MADD): Example simulated trials for inefficient (top) and efficient (bottom) with set-size five are shown. Dashed red and blue lines show decision boundaries for choosing and rejecting an object. The red and blue trajectories represent the decision variable (DV) for good and bad objects. Red and blue patches indicate times subjects viewed a good or a bad fractal, respectively (different shades of blue for different bad objects). In the inefficient search (*θ*=0 top panel), DV for an object was only updated when fixating that object, resulting in serial examination of multiple objects and slow search. In contrast, in the efficient search (*θ*=1 bottom panel), DVs were updated even for peripheral objects, resulting in rapid decision boundary crossing and rapid search. Drift rate (*d*) of non-fixated objects were modulated by *θ* (drift x theta), and there was integration noise sigma (*σ*) added at every time point. (F) Cumulative distribution function (CDF) for drift rate, sigma, and theta in inefficient (orange) and efficient (green) searches. (G) Average search times (solid line) and predicted search times using the MADD model (dashed line) for inefficient (orange) and efficient searches (green) across set sizes in TP trials (left) and TA trials (right). Symbols “=’, “/’, and “X’ indicate significant effects of the main factors: search efficiency, set size, and the interaction between search efficiency and set size, respectively. (H) The scatter plot and correlation between the average theta of a session and the first-saccade target detection rate (up) or search slope (down) across TP trials of that session. Each point is a session with orange and green denoting our binary categorization of inefficient and efficient searches in this plot and similar ones hereafter. The black line is the linear fit in this plot and similar plots hereafter (Demming regression).

As reported previously^10,12^, performance during interspersed choice trials in the value training was high and well above chance for 1-day and +5-day trained fractals (1-day ∼95%, 5+ days 98% accuracies, ts_59_ > 48.6, ps < 2.4e-49 above 50% chance). Nevertheless, search efficiency was different between one-day and +5-day fractals. During the search task, monkeys were tasked with locating good objects among bad ones in target-present (TP) trials or rejecting trials by fixating on the fixation point in target-absent (TA) trials in which all objects were bad (Fig 1D). Trials could have 3, 5, 7, or 9 objects (set-size) displayed on an imaginary circle around the center. The search time slope for TP trials (referred to as the “search slope’) was used to define efficient and inefficient search (see methods). Efficiency was defined by search slopes below the 33rd percentile (<14 ms/item), while inefficiency was indicated by slopes above the 66th percentile (>27 ms/item). While search slope was used to define search efficiency, search time was also significantly shorter in efficient vs inefficient sessions in both TP and TA trials (TP: F_(3, 19192)_ > 167, p < 1e-106; TA: F_(3, 5592)_ > 21, p < 1e-13 Fig 1E; Fig 1E).

### Enhanced Peripheral Object Processing: Behavioral Evidence

Search for a target among multiple distractors can be conceptualized by a <u>m</u>ulti-alternative <u>a</u>ttention-modulated <u>d</u>rift-<u>d</u>iffusion formalism (MADD, also referred to as aDDM^13,14^). In MADD, the accumulation of evidence is strongest for fixated objects and is attenuated by a parameter *θ* < 1 for non-fixated ones which can be taken to model the extent of the peripheral visual processing of objects. The evidence for each object is accumulated by a diffuser until it reaches the selection or rejection bands. The search trial concludes when one diffuser hits the selection boundary, indicating target detection, or when all the diffusers reach the rejection boundary, indicating trial rejection. In addition to *θ*, our model included the drift rate (*d*) and the diffusion noise (*σ*) (see methods). Results showed that MADD was able to fit search times across set sizes and for different search efficiencies (Fig 1E). Examining the parameter fit revealed that the difference in efficient and inefficient search was primarily due to a significant enhancement of *θ* in efficient search (t_137_ = 11.88, p < 1e-25) with minimal effects on drift rate or sigma (*σ* and *d*; t_137_ = 1.3, p = 0.19; t_137_ = −1.18, p = 0.23 respectively, Fig 1F). Indeed, one can verify that *θ* values at extremes of 0 and 1 result in serial (most inefficient) and parallel (most efficient) searches, respectively (Fig 1G). When *θ* = 1, all diffusers rapidly update their values, enabling set-size independent search. In contrast, when *θ* = 0, the target requires multiple saccades since no evidence accumulation can happen for non-fixated objects. Notably, the fitted values of *θ* showed a significant negative correlation with search slopes across all sessions (all sessions, r = −0.70, p < 1e-40 within efficient: r = −0.53, p < 1e-10; within inefficient: r = −0.51, p < 1e-9) and a significant positive correlation with the detecting good target on the first saccade after display onset across sessions (all sessions: r = 0.71, p < 1e-43, within efficient: r = 0.60, p < 1e-14; within inefficient: r = 0.46, p < 1e-8; Fig 1H). In summary, behavioral modeling with MADD revealed an increase in *θ* with search efficiency, suggesting an expansion in the visual processing of peripheral objects in efficient search compared to inefficient search. Next, we explored the neural signature in the vlPFC for this expansion.

### Enhanced Peripheral Object Processing: Neural Evidence

We have previously shown that vlPFC neural response is different in TP vs TA trials within 150ms after the display onset, which can presumably be used to signal the presence of a high-value target in the display (target signal^10^). Importantly, the target signal was found to be stronger in efficient vs inefficient searches across the population (Fig S1A). Given the fact that vlPFC neurons have relatively localized receptive fields^11,15,16^, one expects that the target signal should be a function of the target location (target signal tuning curve). Indeed, such spatial dependence in the target signal is observed in individual neurons (Fig S2) and the population average (Fig S1B). Given such a target signal tuning curve, one can think of at least three scenarios for an overall enhancement of the target signal between inefficient and efficient searches. As shown in Figure 2A, the target signal enhancement can be additive, multiplicative, or involve a spatial widening across angular locations. Any of these tuning curve enhancements can intuitively work to make the neuron more responsive to the presence of a peripheral target across angular locations (Fig S3).

**Figure 2.**
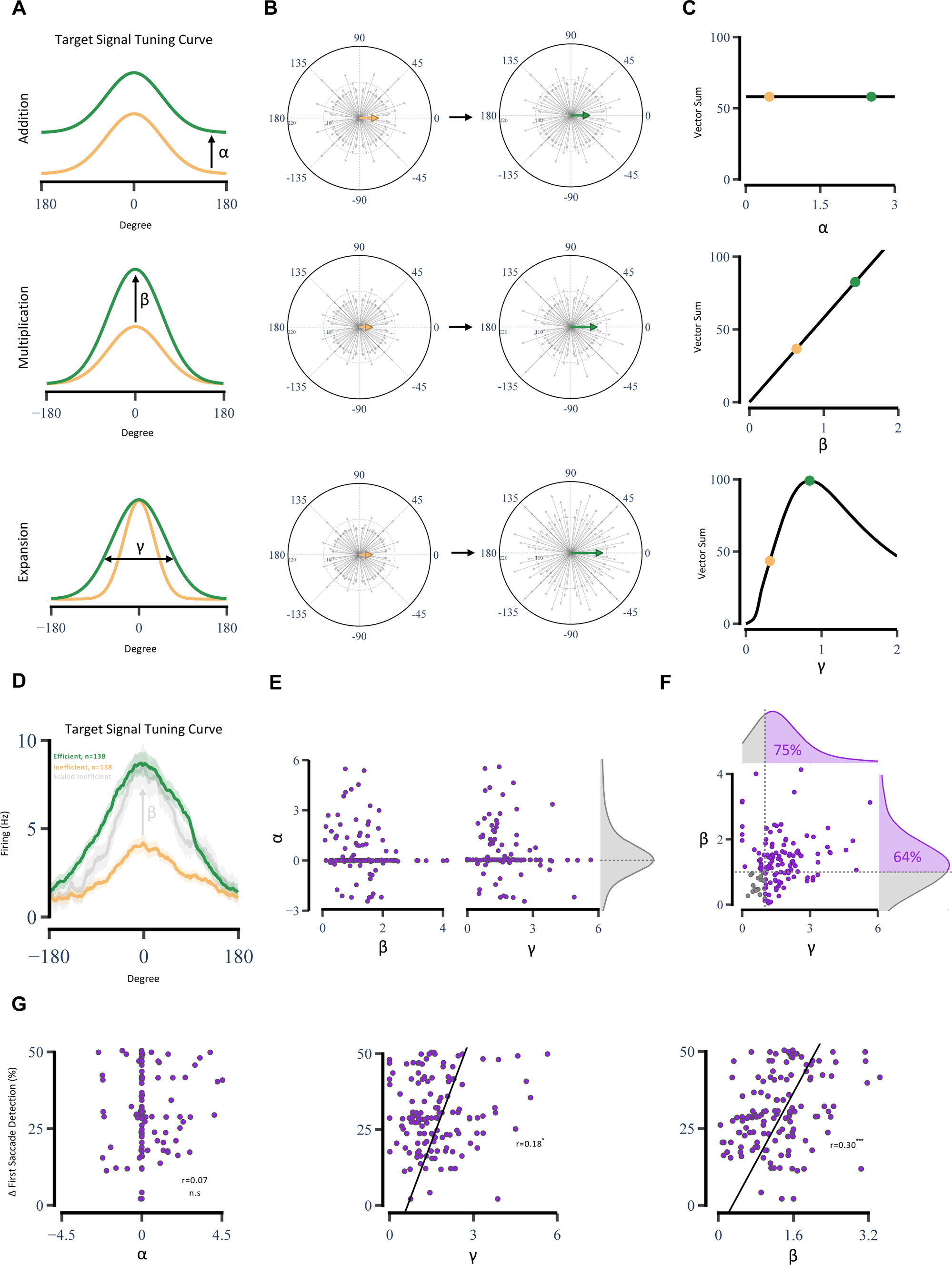
Enhanced spatial processing in vlPFC and its role in target detection. (A) Schematic showing three scenarios for modulation of neural spatial dependent responses to objects (RF tuning curve) assuming object angular location along an eccentric circle around fixation (0 angle being the center of RF). Modulations of the tuning curve by additive (α, first row), multiplicative (β, second row), or expansive (γ, third row) factors are shown. (B) The response of different neurons (10×10 grid) along their preferred direction to an example display (set-size=5) with a target at 0° angle for two different values of each factor. The black arrow shows the population vector sum. Note the larger magnitude for multiplicative and expansive factors but not the additive factor. (C) The effect of each factor on the magnitude of the population vector sum. (D) Population average of target signal tuning curve during inefficient (orange) and efficient (green) searches for neurons with both search types (n=138). The gray trace shows inefficient search tuning curve with multiplicative factor to match the efficient search tuning curve still showing a narrower width. (E) Pairwise scatter plots of additive (α), multiplicative (β), and expansion (γ) factors with marginal distributions. The percentage of neurons with multiplication>1 and expansion>1 are noted. (F) The scatter plot and correlation between each factor and the increase in first saccade target detection in efficient visual search. Note a lack of correlation for the additive factor as predicted in C.

Given that different vlPFC neurons exhibit preferences for various angular locations, a straightforward method to determine the target location would involve calculating the population vector sum of neural activities (Fig 2B)^17–19^. In this model, the probability of a saccade towards a particular direction is assumed to increase as the size of the population vector sum grows larger. Notably, the population vector sum predicts that response multiplication or widening, but not the additive offset, results in a larger resulting vector and, consequently, a higher chance of saccade toward the target (Fig 2C, see methods). Furthermore, while tuning curve multiplication leads to a monotonic increase in the population vector, tuning curve widening initially increases the size of the population vector, followed by a decrease due to the loss of spatial resolution (Fig 2C).

Figure 2D shows the vlPFC population target signal tuning curves for both efficient and inefficient search conditions. These curves are averaged relative to the angular location corresponding to the maximal response (0° corresponding to the maximal point for each neuron; n=138; neurons for both search types, see methods). Interestingly, the efficient search was concurrent with an enhanced tuning curve in vlPFC (for individual neuron examples, see Figure S2). Examining individual neurons revealed a significant percentage of them to show either amplification (β) or widening (γ) or both (One-sample t-test, expected null value=1: t_137_ = 4.55, p < 10e-4; t_137_ = 5.92, p < 10e-7; Fig 2F). Notably, the additive (α) effect was near zero for many neurons (Fig 2G, S5), and the overall effect on the population was not different from zero (t_137_ = 1.88, p = 0.06). Importantly and consistent with our population vector model, the results showed a significant positive correlation between the increase of first saccade target detection in efficient vs. inefficient searches and both the amplification (β) or widening (γ) of value signal. However, no such correlation was found with the additive shift (α) in two-search neurons consistent with the prediction of the population vector sum model (β:r = 0.30, p < 1e-3; γ:r = 0.18, p = 0.03; α:r = 0.07, p = 0.41; Fig 2G).

### Relationship between Target Signal Tuning Curve, *θ*, and Search Efficiency

In order to use a single metric that combines target signal enhancement across angular locations, we used the <u>a</u>rea <u>u</u>nder the target signal tuning curve (AUT, Fig 2D). We could confirm a significant rightward shift in the cumulative density function (CDF) for efficient search across the vlPFC population (t_137_ = 9.22, p < 1e-15, Fig 3A). The enlargement of the RF size was also confirmed for neurons with both search types (Fig 3B).

**Figure 3.**
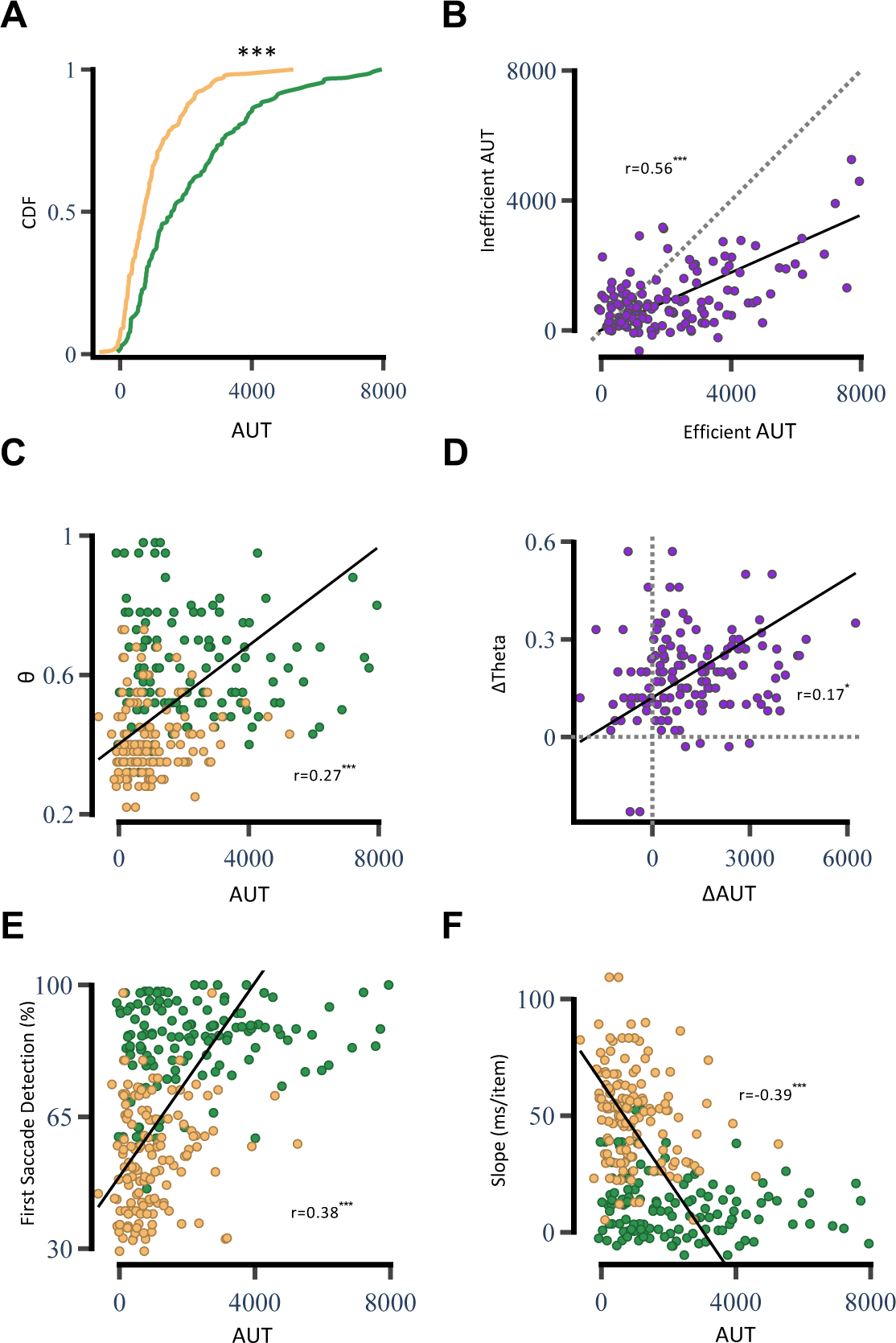
Relationship between spatial processing in vlPFC, MADD *θ*, and search efficiency. (A) Cumulative distribution (CDF) of the area under target signal tuning curve (AUT) during efficient (green) and inefficient (orange) searches in neurons with both search types (n=138). (B) Scatter plot of AUT in efficient vs inefficient searches in neurons with both search types (n=138). (C) Scatter plot and correlation between *θ* and AUT across all sessions. (D) Scatter plot and correlation between change in *θ* and change in AUT from inefficient to efficient search for neurons with both search types. (E) Scatter plot and correlation between first saccade detection of target and AUT. (F) same as E but for TP search slope and AUT.

Behavioral fits using MADD suggested the increase in *θ* to be the underlying mechanism for search efficiency (Fig 1F). Neural analysis revealed that the enhancement of AUT was concurrent with efficient search (Fig 3A-B). This naturally begs a question about the relationship between the *θ* and the AUT. Results showed a significant and positive correlation between the AUT of the neuron recorded in a session and the *θ* of that session (Fig 3C, r = 0.27, p < 1e-5). Once again, in the neurons that were recorded in both efficient and inefficient search blocks, the increase in AUT was also found to be correlated with the increase in *θ* when comparing inefficient to efficient search (Fig 3D, r = 0.17, p < 0.04). The increase in AUT was also correlated with measures of search efficiency such as higher first saccade detection (r = 0.38, p < 1e-10, Fig 3E) and lower search slopes (r = −0.39, p < 1e-10, Fig 3F). It is important to note that the correlation between the responses of a single neuron is expected to be an inherently noisy correlate of search accuracy compared to *θ*, which models the behavior orchestrated by a large number of neurons. Thus, observing these significant correlations between single neuron measures and behavior is notable.

## Discussion

Visual search for targets not distinct from their surroundings by low-level guiding features is thought to be set-size dependent and time-consuming^1,2^. Search becomes especially unforgiving when the number of targets to look for increases, possibly due to limited working memory capacity^20–22^. Nevertheless, we have recently shown that valuable objects can be found efficiently in the visual search despite their large number and lack of low-level guiding features^10,12,23^. While low-level pop-out is believed to arise from parallel processing of color, orientation, and size across the visual field in early sensory areas^24–26^, a similar mechanism for valuable objects with arbitrary shapes is unknown. Here, we aimed to provide a neural mechanism for valuable object pop-out by analyzing vlPFC neuron activity, which is known to be involved in visual search^27,28^ and value memory^15^ along with behavioral modeling using MADD. Our behavioral analysis revealed efficient search arises from enhanced spatial processing for valuable objects in the periphery (Fig 1). The neural data showed amplification and spatial widening of the vlPFC target signal tuning curve (Fig 2). Amplification and widening of the target signal tuning curve were predicted to increase the detection of the target by first saccade using a population vector sum model and were subsequently confirmed to be correlated with behavioral measures of search efficiency (Fig 2-3). Together, these data suggest enhancement of the target signal tuning curve in vlPFC to underly value-driven search efficiency.

Visual search for a target among other objects can be framed as a decision-making task in which the subject has to evaluate multiple objects, decide which one is the target, and reject others. We formulated this decision-making problem using a multi-alternative attention-modulated drift-diffusion (MADD) model. Variants of this model have been previously used to fit search times in visual search^13,14^. In particular, our model assumes that evidence accumulation is fastest for foveated objects but is attenuated by a *θ* parameter for peripheral objects (0 < *θ* < 1, hence the attentional modulation). Behavioral fits revealed that efficient search was concurrent with smaller attenuation (higher *θ*) for evidence accumulation of peripheral objects but with minimal changes in drift rate (*d*) or integration noise (*σ*) (Fig 1F). The increase in *θ* indicates a move toward simultaneous evaluation of objects regardless of locus of gaze and suggests such simultaneity as the underlying cause for value-driven efficient search.

vlPFC was already implicated in spatial and feature-based visual search^28–30^. We have recently shown that vlPFC encodes efficiency in value-driven visual search independent of low-level guiding features as well^10^. Here, we have expanded our previous findings by showing that the spatial tuning of the target signal in vlPFC becomes magnified and wider across angular locations during efficient search. We showed that both of these effects, but not a simple additive shift in the target signal tuning curve, predict an increase in first saccade target detection by a simple population vector sum method (Fig 2). Furthermore, the area under the target signal tuning curve was significantly correlated with the *θ* parameter in MADD (Fig 3). These findings suggest that the target signal in vlPFC could be the neural correlate for *θ* in MADD. However, we do not observe the neural correlate of the evidence accumulation or drift rate (*d*) in vlPFC (Fig S1). One possibility is that evidence accumulation could be done by areas receiving the vlPFC inputs, such as in SC^31^ or LIP^32,33^. We have recently shown neural correlates of evidence accumulation in value-driven food choice using simultaneous fMRI-EEG data in the insular cortex in humans^34^. Similar whole-brain explorations in macaques would be needed to find neural correlates for MADD components and confirm its validity as a suitable model of visual search.

While the mechanism for enhancing spatial tuning in our task is unknown, it will likely involve reward-related plasticity within the cortico-basal ganglia loop where vlPFC participates^35,36^. Indeed, our recent work has shown an enhancement of the target signal in Substantia Nigra reticulata (SNr)^23^, which receives indirect input from vlPFC and also projects back to it via thalamus^37^. It would be interesting to investigate whether reward-dependent synaptic plasticity, enabling vlPFC neurons to integrate inputs across a larger pool, could result in a wider receptive field for valuable objects. Considering the role of dopamine in encoding value learning and memory^38^, as well as in synaptic plasticity in the cortex and striatum^39^, it remains to be determined whether this response enhancement involves dopamine-dependent synaptic plasticity.

In our task, the number of potential targets in a given session was large (>12), and targets were not cued but instead were singled out by past reward history. This type of visual search differs from well-studied searches in which a cued target is seen immediately before display onset, which are known to rely on working memory^22^ and top-down priority maps^40^. Our value-driven search is not consistent with a slew of bottom-up salience maps^41^ due to a lack of guiding features and random object-reward associations. Instead, our results provide evidence for a third category of salience different from top-down and bottom-up controls, as alluded to previously^42^. We refer to this as memory-based salience. Indeed, our recent work has implicated vlPFC in other aspects of memory-based salience involving aversive associations and perceptual novelty/familiarity^43^. Evidence shows that visual attention enhances the gain of the spatial tuning curve of neurons in visual processing areas such as middle temporal (MT) and V4^44–46^. This prompts the possibility that a similar memory-based attentional mechanism may underlie the receptive tuning curve enhancement of the target signal observed in vlPFC neurons.

In summary, our findings put forth a conceptual and neural framework for understanding valuable object pop-out during visual search. Our results demonstrate that long-term value association results in amplification and spatial widening of the target signal tuning curve in vlPFC neurons, thereby enhancing the spatial processing of peripherally presented valuable objects, consistent with behavioral modeling using MADD. More broadly, our findings are a striking example of how value learning can change the neural tuning of PFC neurons for fast detection of valuable objects. These results point to value-driven plasticity within vlPFC or its input areas to aid in the detection of valuable objects. The specifics of synaptic and network mechanisms underlying such reward-related enhancements in neural responses remain to be investigated.

## Material and Methods

### Subjects and Surgery

Two male rhesus macaques, monkeys S (12 years, 10kg) and H (10 years, 12kg), were used in this study. All methods and animal care were approved by the Ethical Committee of the Institute for Research in Fundamental Sciences (IPM) and adhered to the standards established by the National Institutes of Health (USA) for the Ethical Treatment and Use of Laboratory Animals (IPM, protocol number 99/60/1/172). Initially, a titanium head holder and a recording chamber were surgically implanted on the heads of each animal under general anesthesia. The head holder was positioned along the midline of the parietal lobe for both monkeys. Additionally, a recording chamber was placed on the right prefrontal cortex (PFC) for Monkey S and on the left PFC for Monkey H, with a lateral tilt. To verify the accurate placement of the recording chamber post-surgery, MR images of monkeys’ heads were acquired. After the monkeys had become proficient in the experimental tasks, a second surgery involving a craniotomy over the vlPFC area was conducted. Recording neuronal data was done using a 1-mm-diameter grid that fitted inside the chamber. Parts of the data in this work were previously reported by Abbaszadeh et al.^14^, which also provides further details on recording location, stimuli, and tasks used.

### Recording Localization

T1- and T2-weighted MR images (3T, Prisma S5iemens) were used to map precise recording locations within the ventrolateral prefrontal cortex (vlPFC). To further validate the positional accuracy of each animal’s vlPFC, the National Institute of Mental Health Macaque Template (NMT) toolbox was used to morph the standard monkey brain atlas with the native space of the monkeys^47,48^.

### Stimuli

Fractal-shaped objects^49^, as shown in Figure 1A, were employed as visual cues. Each of these fractals featured a shared core encircled by four point-symmetrical polygons, superimposed with smaller ones at the forefront. The attributes of each polygon (such as size, edges, and color) were randomly selected. The average diameter of the fractals was 4 degrees. Monkeys were exposed to numerous fractals (over 600) across various tasks. For the search task, monkeys S and H each saw 736 and 824 fractals, respectively. The monkeys viewed these fractals either during a single training session or throughout five reward training sessions, resulting in a collection of “over-trained” fractals.

### Task Control and Neural Recording

A custom software program developed in the C language was employed to manage both behavioral activities and recordings. A Cerebus Blackrock Microsystem was used to collect neural data (www.blackrockneurotech.com). Eye tracking was conducted using Eyelink 1000 Plus, operating at a sample rate of 1000 Hz. During each trial, the reward consisted of apple juice diluted with water (50% and 60% dilution for monkeys S and H, respectively). A total of 526 well-isolated, visually sensitive neurons were recorded for this study, primarily concentrated in area 46v, situated ventral to the principal sulcus (230 from Monkey S and 296 from Monkey H). These neurons were classified based on the search slope criteria described in the results section. They were recorded during either efficient, inefficient, or both types of searches, with 118 and 177 neurons in the efficient search and 122 and 178 in the inefficient search for monkeys S and H, respectively. In total, 138 out of 525 neurons were recorded in both efficient and inefficient searches.

### Object-value Training Task

A biased value saccade task was employed to train the object values in monkeys (Fig. 1C). During each task trial; the monkey was directed to fixate on a white dot appearing at the center of the screen. Upon maintaining fixation for 200ms, a high-value (good) or low-value (bad), fractal was displayed at one of eight peripheral locations (eccentricity of 9.5°). Following a 400ms overlap period, the central fixation point disappeared, prompting the animal to execute a saccade towards the fractal and sustain gaze fixation for an additional 500 ± 100ms to obtain either a large (for good fractal) or a small (for bad fractal) reward. After reward delivery, a variable inter-trial interval (ITI) ranging from 1 to 1.5 seconds ensued, during which a blank screen was presented.

Each training session encompassed 80 trials, comprising 64 force trials wherein each object was presented eight times in a pseudo-random order and 16 choice trials wherein one good and one bad fractal were simultaneously displayed in a diametrical arrangement on the screen. The timing structure for the choice trials was the same as the force trials, with the distinction that the monkey was required to select one of the fractals with a saccade. Successful execution of each trial triggered a correct tone. In contrast, an error tone was triggered in cases of early saccade to a fractal or a disruption of fixation. The outcomes observed during choice trials were used to quantify object value learning in each monkey.

### Value-driven Search Task

In this task, monkeys were required to identify a good object (target) within a variable number of bad objects during target-present (TP) trials or to reject the trial by returning to the center during target-absent (TA) trials, where all presented objects were bad^12^. TP and TA trials were intermixed with equal likelihood. Monkeys learned object values using object value training tasks prior to the search. The start of each trial was signaled by a fixation dot displayed on the screen (Fig 1D). After a 400-millisecond of central fixation, sets of 3, 5, 7, or 9 fractals were presented in an imaginary circle, equally spaced, with a 9.5° eccentricity (display onset). The angular location of the target was taken from a uniform distribution around the circle. Objects were pseudo-randomly selected from a set of 24 fractals (12 good and 12 bad) for each trial, depending on the trial type (TP or TA). Following the display onset, monkeys had three seconds to either choose an object by fixating it for a minimum of 600ms (committing time), followed by an additional 100 ms for reward receipt, or to reject the trial by remaining at the center (for 900ms) or by returning to central fixation and staying for 300ms after making saccades. Monkeys were permitted to make as many saccades as they wished during the three seconds and to shift their gaze away from the object before the committing time elapsed (gaze breaks during the subsequent 100ms would result in an incorrect tone). Monkeys would receive a high reward for choosing good and a low reward for choosing bad objects. Rejecting a trial led to quick progression to the subsequent trial (after 400ms), with a 50% chance of encountering a good object in a target-present trial. For non-rejected trials, a randomized inter-trial interval (ITI) of 1 to 1.5 seconds was implemented after reward delivery. Monkeys would also receive an error tone if no object was chosen after three seconds and the trial was not rejected either (rare, <0.1% of trials).

### Search Type Categorization

The search sessions were classified as efficient or inefficient based on the search slope, calculated using linear regression (scikit-learn v1.3.1) for TP trial search times across different set sizes. Sessions with a search slope below the 33rd percentile of the search slope distribution were designated efficient, and sessions with a search slope above the 66th percentile were categorized as inefficient.

### Behavioral Data Analysis: Multi-Alternative Attention-Modulated Drift Diffusion Model (MADD)

We used a drift-diffusion framework to model the decision-making process during the search task. In this model, each object has its own decision variable or diffuser. Thus, a given trial, depending on the set size, had diffusers ranging from 3 to 9. Each diffuser started accumulating evidence for its corresponding fractal being the search target or not from the display onset. A decision involved either selecting a fractal as the target or rejecting the entire trial as the target-absent with absorbing decision boundaries. Hitting the upper boundary signified acceptance of the corresponding fractal as the target. Conversely, if a diffuser reached the lower boundary, it marked its fractal as a bad object. In this scheme, trial rejection happened if and when all diffusers hit the lower boundary (Fig1G). Model parameters included drift rate (*d*), integration noise standard deviation (*σ*), and the attention modulation ratio for non-fixated over fixated objects (*θ*). These parameters were fit separately based on trial type (TP/TA), set size, and search type (efficient/inefficient).

The diffuser’s value at each time step is calculated using:

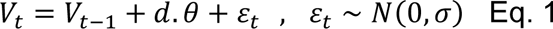

Here, “V’ represents the decision variable, “t’ represents the time of a single trial from 0 to 3000ms (max search time), and “*ε*’ represents the diffuser noise. The attention modulation ratio is defined as:

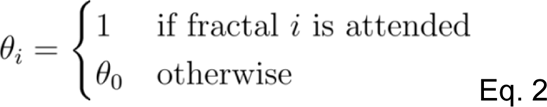

The idea here is that the decision variable can accumulate information even when the fractal is in the subject’s periphery. When the subject fixates on a fractal, the decision variable updates as usual with the rate of *d*. At the same time, the decision variables of other fractals may update in proportion to the *θ*_0_ ratio. In the extremes, when *θ*_0_ is set to 0, objects have to be sequentially fixated and processed, but when *θ*_0_ set to 1, all objects are processed in parallel without the need to fixate them.

To estimate model parameters for different search task conditions within each search run, we utilized the grid search method to identify optimal values. The mean squared error (MSE) served as the cost function to determine optimal values for each condition. The model’s prediction was the search time (i.e., time to locate the target in TP or reject trial in TA trials). For a given trial, we calculated the prediction for search time by averaging the values obtained from 100 iterations of the drift-diffusion process. Considering the range of search time variations observed in monkey behavior and model predictions, we set valid parameter ranges to be [0, 1] for *θ*, [0, 0.1] for *σ*, and [0, 0.01] for d.

### Population Vector Sum Model

We simulated the effect of target signal tuning curve size on detecting the target by first saccade by tiling the visual scene with a uniform grid of 10×10 neurons. Each neuron spatial RF was modeled as a bivariate Gaussian distribution whose value at a given location signified the neuron’s firing to an object appearing at that location. The response range for each neuron was chosen to match the values observed in our vlPFC population of neurons with a minimum (fr_min_=16Hz) and maximum (fr_max_=41Hz) during the 100-400ms after display onset. A TP trial was simulated by putting a variable number of fractals (3, 5, 7, or 9) on a circle around the center of the screen. The fractal at 0 degree was the good object (target).

We first calculated the activity of each neuron depending on the location of objects and the set size using the following equation:

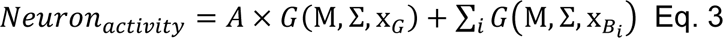

*A* is the ratio of activity to good vs bad (default 1.4)

*G*(Μ, Σ, x) is the neuron’s RF, which is a bivariate Gaussian distribution centered at Μ (on the 10×10 grid) with symmetric variance Σ, evaluated at location x

*x*_*G*_ location of good object

*x_B_i__* location of i^th^ bad object

We then multiplied this activity by a vector that started from the center of the screen and pointed to the center of the neuron’s RF (i.e. Μ). We then calculated the vector sum across the 10×10 grid of neurons. The direction of this vector pointed to the direction of the saccade (which would always be toward a good object due to A>1), and its magnitude was assumed to be monotonically related to the probability of saccade to that direction (an increase in vector magnitude means higher saccade probability).

We considered three scenarios for modulations of neurons RF, including additive (α), multiplicative (β), and expansive (γ) mechanisms. The parameter α accounts for baseline shifts, the parameter β accounts for gain, and the parameter γ accounts for changing the width of the bivariate Gaussian, respectively. The 2D target signal is defined as the difference in firing to TP minus TA with matched set size and object locations. Note that the subtraction of these two entities gives us the same bivariate Gaussian distribution as the target signal for that neuron, and thus, we considered the same modulations on the real neuron’s target signal in the following section.

### vlPFC Target Signal Tuning Curve

To quantify the target signal dependence on the angular location of the target, we first calculated the neuron response (100-400ms epoch after display onset) as a function of the target’s angle in TP trials. Next, we applied a circular moving average (window length=30°) to each neuron’s angular response to the target location and performed linear interpolation with a 1° resolution. This allowed us to arrive at the TP tuning curve for each neuron with 1° resolution (360 data points). To calculate the target signal, we subtracted the average response for TA trials from this TP tuning curve for TP-preferred neurons 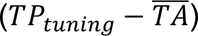. In TA-preferred neurons, this subtraction was performed reversely 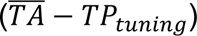. Here,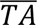 is a scalar equal to the average response collapses across all TA trials. We call the result of this subtraction the *uncorrected* target signal tuning curve (Fig S4A; for reasons below). The peaks of these tuning curves were rotated to zero to align them and then averaged across the population.

Since averaging the target signal tuning curve across neurons aligned to their peaks can create a false average peak even without any spatial tuning, we applied the same technique to the TA trials as a control (Fig S4B). In TA trials, we treated the first fractal as a pseudo-target to calculate the tuning curve for TA trials with a 1° resolution using the same method as above. We then created a *fake* target signal by doing 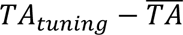 for TP-preferring neuron and 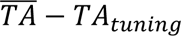 for TA-preferring neurons. The peaks of this fake target signal were also rotated to zero and were averaged across the population. We then subtracted this average *fake* target signal from the average uncorrected target signal (*uncorrected-fake*) to arrive at the final target signal tuning curve shown in Fig 2D. The area under this tuning curve is termed the AUT and is used in Fig 3.

To calculate the modulation of target signal between efficient and inefficient search, for each neuron, we fitted α, β, and γ ratios (Figs 2E-F) by using the Nelder-Mead algorithm within the Python SciPy package using Eq. 4:

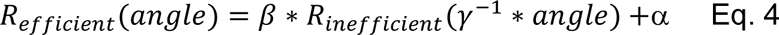

where *R* is the target signal circular tuning curve.

Note that while α and β in the bivariate Gaussian simulation will be equal to those estimated in Eq. 4, the *γ* fitted in Eq. 4 will not be the same as the one used for simulating the effect of bivariate Gaussian since we are measuring responses on a circle and and not on the whole 2D grid. Nevertheless, the fitted *γ* will be monotonically related to the 2D tuning curve *γ* (i.e., an increase in the width of bivariate Gaussian will not decrease the width of the circular tuning curve).

### Statistical Tests and Regressions

A one-way ANOVA test was utilized to examine the significant impact of set size on search time. In addition, a two-way ANOVA test was employed to explore the combined influence of search efficiency and set size on these same variables (Fig 1E). For the linear line fit, we used type II (Demming) Regressions^20^ (Figs 1H, 2G, 3B-F). Pearson’s correlation coefficient (r) was used to measure correlations in all figures. Error bars or shades in all plots show the standard error of the mean (SEM). The significance threshold for all tests in this study was p < 0.05. ns: not significant, *p < 0.05, **p <0.01, ***p < 0.001 (two-sided).

## Declaration of interests

The authors declare no competing interests.

---

**Figure S1.**
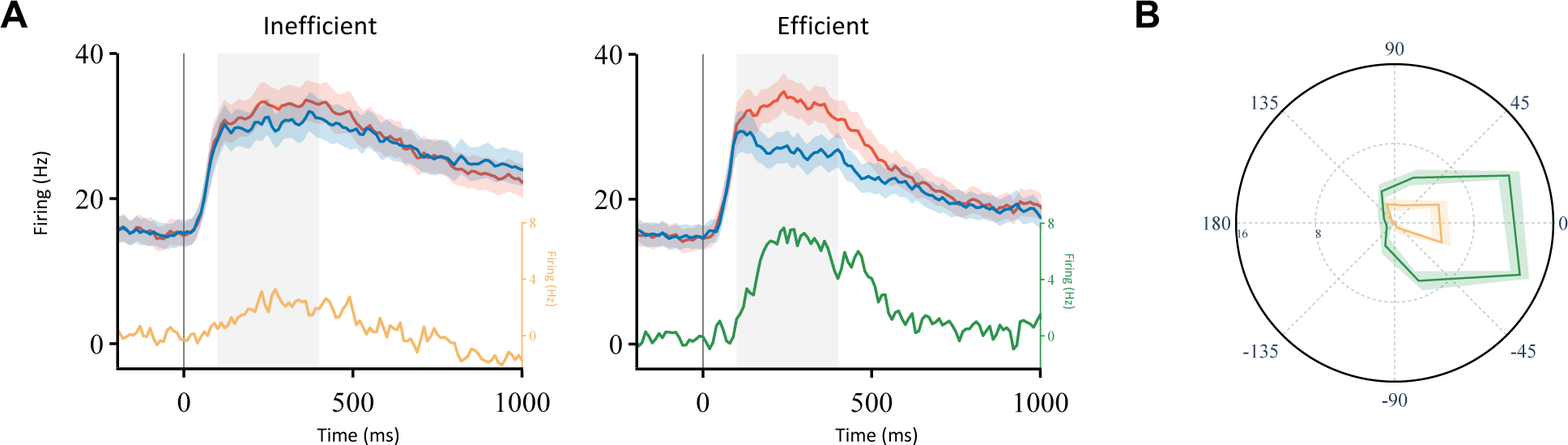
Firing rate and target signal in inefficient and efficient searches. (A) Population average of peristimulus time histograms (PSTH) for two-search neurons (n=138) during efficient (right) and inefficient (left) searches, illustrating TP trials (red), TA trials (blue), and the subtraction of TP and TA trials representing the target signal (in green for efficient and in orange for inefficient). (B) The polar plot shows the average target signal of two-search neurons (binned at eight radial directions) in both inefficient (orange) and efficient (green) searches. The peaks of these target signals were rotated to zero to align them before averaging over population (uncorrected target signal).

**Figure S2.**
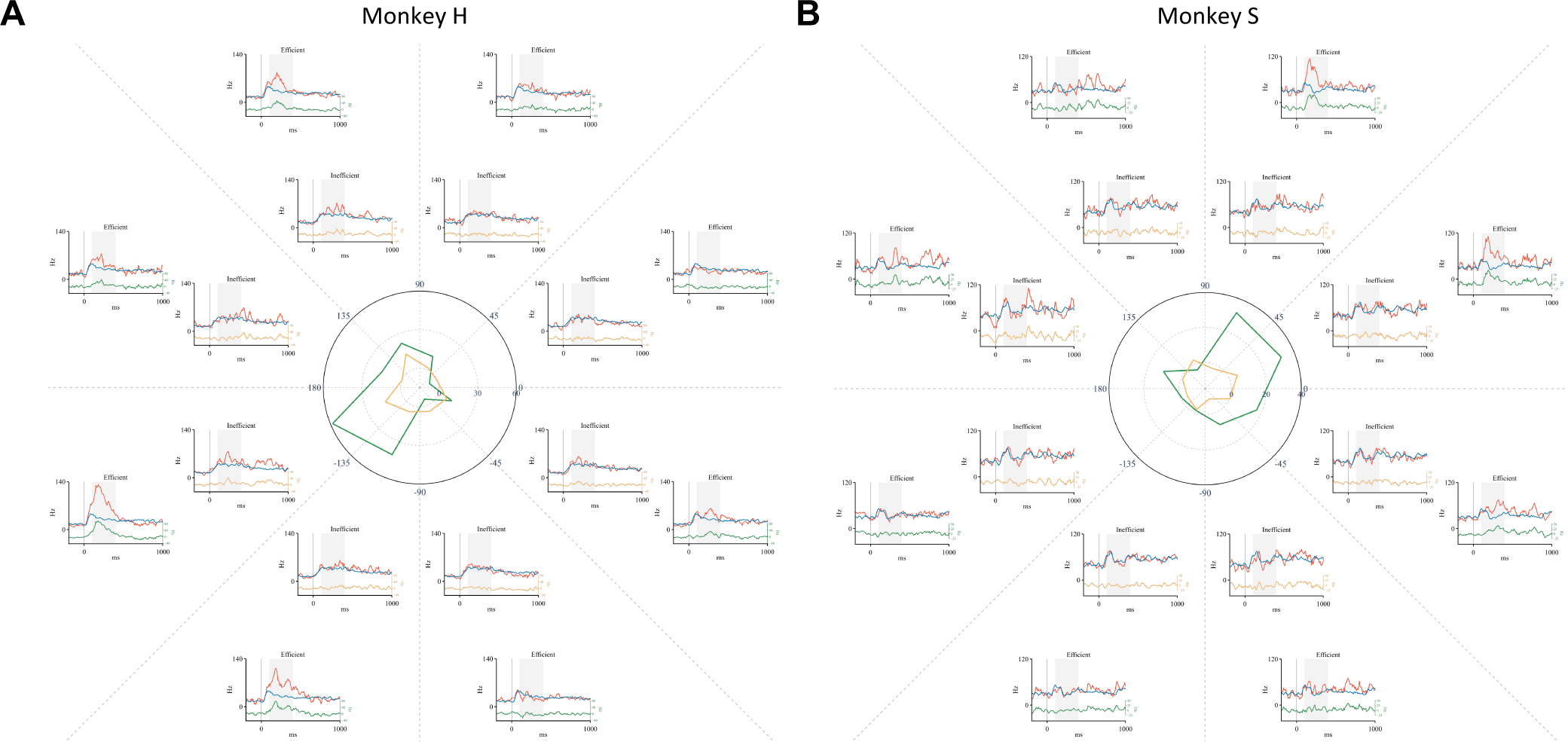
Example neurons and their target signal tuning curves. The polar plot shows the target signal of the example neuron from each monkey across various target locations (binned at eight radial directions) in both inefficient (orange) and efficient (green) searches. Each of the eight sections in the plot displays the PSTH of the neuron for inefficient search (inner plot) and efficient search (outer plot), separately for TP trials (in red), TA trials (in blue), and the subtraction of TP and TA trials representing the target signal (in green for efficient and in orange for inefficient).

**Figure S3.**
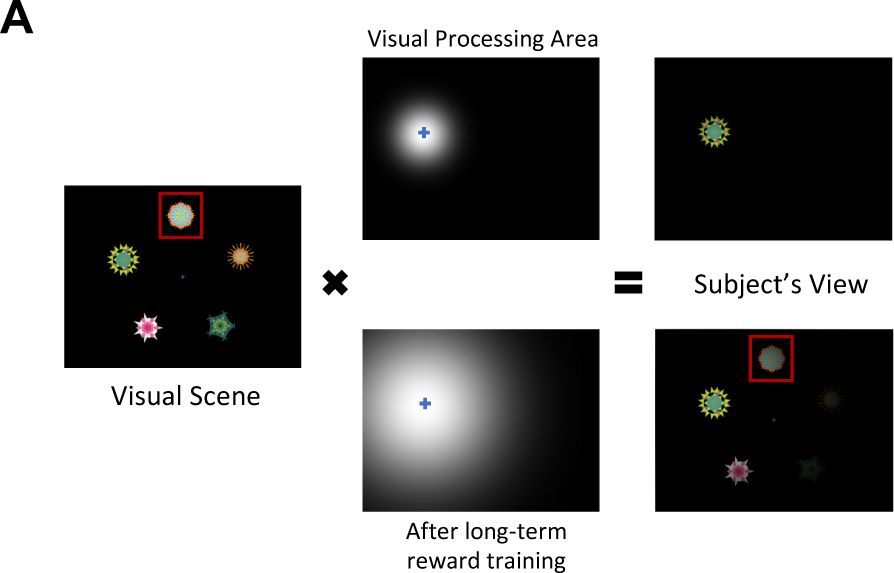
Schematic showing how enlargement of neuron spatial processing allows it to see the target within its RF. Example TP trial shows set-size 5 with a target on top of the screen (left column). The neuron RF is modeled as a bivariate Gaussian centered at a peripheral location (middle column top row). The RF width got larger due to object reward training (middle column bottom row). The RF is multiplied by the scene to show the objects affecting the neuron’s firing (right column). The target becomes visible to the neuron in the enlarged RF condition.

**Figure S4.**
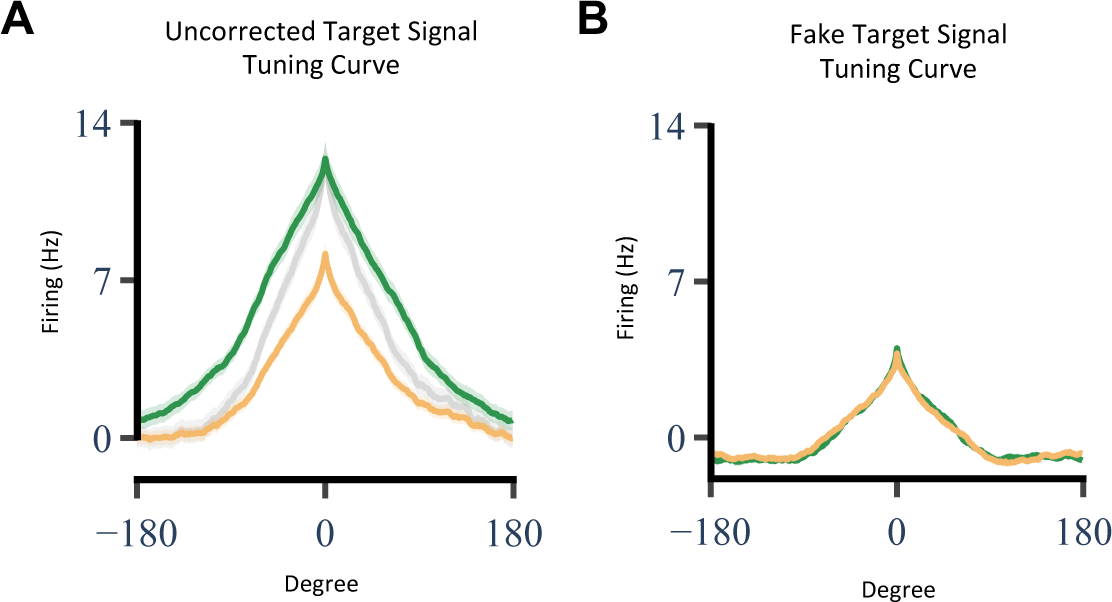
Correction of target signal tuning curve. (A) Uncorrected and (B) Fake target signal tuning curve, with details comparable to those described in Fig 2D (see methods).

## Notes

### Competing Interest Statement

The authors have declared no competing interest.

